# GO-PCA: An Unsupervised Method to Explore Biological Heterogeneity Based on Gene Expression and Prior Knowledge

**DOI:** 10.1101/018705

**Authors:** Florian Wagner

**Affiliations:** PhD Program in Computational Biology and Bioinformatics, Duke University

## Abstract

Genome-wide expression profiling is a cost-efficient and widely used method to characterize heterogeneous populations of cells, tissues, biopsies, or other biological specimen. The exploratory analysis of such datasets typically relies on generic unsupervised methods, e.g. principal component analysis or hierarchical clustering. However, generic methods fail to exploit the significant amount of knowledge that exists about the molecular functions of genes. Here, I introduce GO-PCA, an unsupervised method that incorporates prior knowledge about gene functions in the form of gene ontology (GO) annotations. GO-PCA aims to discover and represent biological heterogeneity along all major axes of variation in a given dataset, while suppressing heterogeneity due to technical biases. To this end, GO-PCA combines principal component analysis (PCA) with nonparametric GO enrichment analysis, and uses the results to generate expression signatures based on small sets of functionally related genes. I first applied GO-PCA to expression data from diverse lineages of the human hematopoietic system, and obtained a small set of signatures that captured known cell characteristics for most lineages. I then applied the method to expression profiles of glioblastoma (GBM) tumor biopsies, and obtained signatures that were strongly associated with multiple previously described GBM subtypes. Surprisingly, GO-PCA discovered a cell cycle-related signature that exhibited significant differences between the Proneural and the prognostically favorable GBM CpG Island Methylator (G-CIMP) subtypes, suggesting that the G-CIMP subtype is characterized in part by lower mitotic activity. Previous expression-based classifications have failed to separate these subtypes, demonstrating that GO-PCA can detect heterogeneity that is missed by other methods. My results show that GO-PCA is a powerful and versatile expression-based method that facilitates exploration of large-scale expression data, without requiring additional types of experimental data. The low-dimensional representation generated by GO-PCA lends itself to interpretation, hypothesis generation, and further analysis.

## Introduction

Genome-wide expression profiling, or *transcriptomics*, is a highly popular approach for obtaining a systematic view of the molecular heterogeneity underlying samples of cells, tissues, tumor biopsies or other biological specimen. The success of transcriptomics is based on microarray and high-throuhgput sequencing technologies, which have led to reductions in costs and improved measurement accuracies. Currently, the development of single-cell transcriptomics methods is promising a dramatic increase in resolution, enabling systematic explorations of heterogeneity at a level of detail never seen before [1] (see e.g. [2, 3], recent applications of single-cell transcriptomics in the fields of developmental biology and cancer research, respectively).

Considering the pace with which large-scale transcriptomic datasets can be produced, data analysis often represents a significant bottleneck. The machine learning literature offers a plethora of unsupervised methods which have been adopted to various degrees for the exploratory analysis of gene expression data. Popular approaches include principal component analysis (PCA) [4], hierarchical clustering [5], k-means clustering, consensus clustering [6], non-negative matrix factorization (reviewed in [7]), mixture models (e.g., [8]), and many others. These methods can be characterized as *generic*, in that they operate based on general principles (e.g., prinicipal components are uncorrelated and capture maximum amounts of variance), and do not take any specific biological aspects of the data into account.

While applications of the aforementioned methods have led to profound insights into biological processes (e.g., the identification of clinically relevant cancer subtypes [9, 10]), arriving at such results typically requires significant human effort combined with expert knowledge, and can be fraught with difficulties. In many cases, the data contain significant but unknown biases which can obscure interesting signals and create spurious results (e.g., batch effects [11]). Furthermore, the output of unsupervised methods often consists of clusters or factors containing hundreds of genes, which are difficult to interpret and necessitate further analysis before any biological intuition can be applied.

These challenges motivate the development of more specialized tools for the exploratory analysis of transcriptomic data that 1) improve the detection of biologically relevant patterns, 2) confer robustness with respect to unkonwn biases and batch effects, and 3) yield readily interpretable results that facilitate hypothesis generation. The incorporation of *prior knowledge* into unsupervised algorithms provides a major opportunity for achieving these goals. In principle, prior knowledge can bias the analysis in favor of biologically plausible results, thereby reducing the influence of extraneous biases such as batch effects, which do not exhibit biologically meaningful patterns. It can also help provide meaningful labels for discovered patterns, which in turn facilitates the interpretation of results [12].

In light of the intuitive appeal of this idea, as well as its highly successful application in supervised settings [13], there exist surprisingly few methods that exploit prior biological knowledge in a general unsupervised setting. Several methods have been designed for the narrow task of identifying regulatory relationships ([14] and ref. 11-14 in [12]). For more general purposes, it has been proposed to adjust the distance metric used in hierarchical clustering by a term that quantifies similarity of GO or KEGG annotations between pairs of genes, with a tuning parameter allowing for a flexible trade-off between knowledge-based and data-driven analysis [15, 16]. Annotation-based adjustments have also been proposed for use in k-means/k-medioid clustering [17, 18, 19] and mixture models [20].

The method proposed here relies on PCA, one of the most versatile unsupervised methods, and takes a novel approach for integrating prior knowledge. Rather than adjusting an internal metric, the method synergistically combines PCA with nonparametric GO enrichment analysis, in order to define expression signatures based on compact sets of correlated and functionally related genes. In this way, the method aims to produce an easily interpretable, low-dimensional representation of biologically relevant patterns that are associated with diverse axes of variation.

## Results

### GO-PCA combines principal component analysis (PCA) with nonparametric GO enrichment analysis

In order to facilitate exploration of transcriptomic data, I sought to design a method that would be able to analyze all major axes of variation, identify functionally related sets of genes that showed high correlation along each axis, and present the results in an easily interpretable fashion. To this end, I developed the GO-PCA algorithm, named after its two building blocks, PCA [4], and GO enrichment analysis based on gene annotations from the UniProt-GOA database [21]. GO-PCA first performs PCA on the dataset (Figure 1a), and then tests each principal component (PC) for enrichment of functionally related genes. More formally, for each PC, genes are ranked by their loadings, and a p-value based on the *minimum hypergeometric* (mHG) test statistic [22] is calculated for each of about 4,000 GO terms (Figure 1b). Then, a stringent Bonferroni correction is applied to the p-values, and the genes associated with each enriched term are used to construct an expression signature. GO-PCA excludes GO terms that are either too broad or too specific, and prioritizes and filters enrichments based on their estimated effect size (fold enrichment), in order to avoid redundancies that arise from closely related GO terms. The methodology is described in detail in the Methods section.

**Figure 1.**
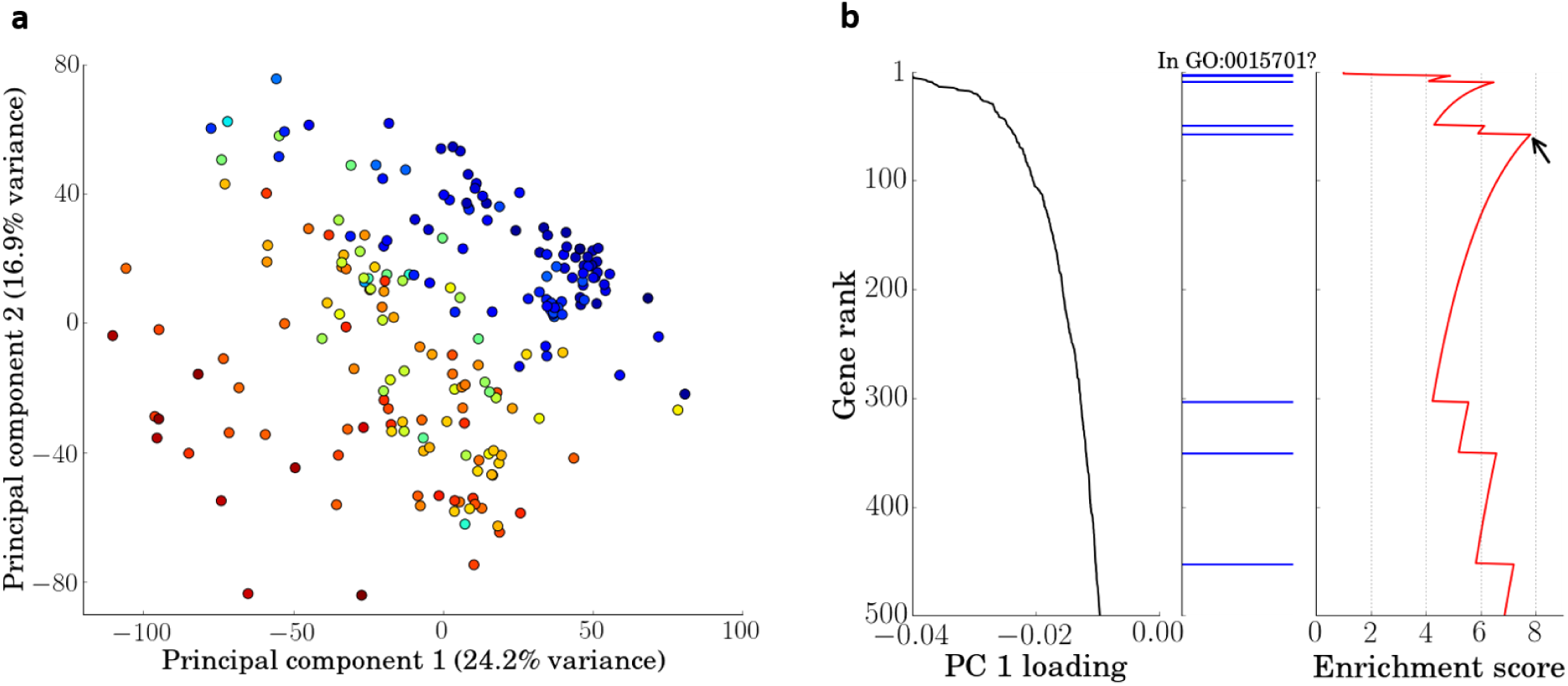
Building blocks of GO-PCA. **a** PCA is performed on the gene expression matrix, treating the genes as variables and the samples as observations. Shown are the principal component (PC) scores of the 211 samples in the hematopoietic dataset [23], for the first two PCs. Color codings correspond to the expression level of the gene *HBA1* (Hemoglobin subunit alpha; red = high, blue = low), which contributes heavily to PC 1. **b** For each PC, genes are ranked by their loadings (left; only the first 500 genes are shown). The ranked list of genes is used to test for enrichment of approx. 4,400 GO terms. The middle graph shows the ranks of genes associated with the GO term “bicarbonate transport” (GO:0015701) in the list, and the graph on the right shows the enrichment score calculated at each threshold by the minimum hypergeometric (mHG) test [22], in order to find the threshold with the best score (black arrow). The mHG algorithm then calculates an exact p-value for this score.

### Application of GO-PCA to a diverse panel of cell populations from the hematopoietic lineage recovers many cell-type specific characteristics

As a first test of my method, I aimed to assess whether GO-PCA, upon presentation with highly heterogeneous transcriptomic data, would be able to produce biologically meaningful signatures that were both functionally diverse and easily interpretable. I therefore applied GO-PCA to a dataset comprising 211 transcriptomes that represented 14 different hematopoietic lineages [23]. GO-PCA produced 43 signatures containing between 5 and 39 genes (Figure 2). The signatures were derived from GO terms representing diverse biological processes, including DNA replication, mitochondrion degradation, and various immune system-related processes. Although some signatures were highly correlated, there were multiple clearly distinguishable clusters, as well as complex patterns of partial correlation and anti-correlation (Figure S1).

**Figure 2.**
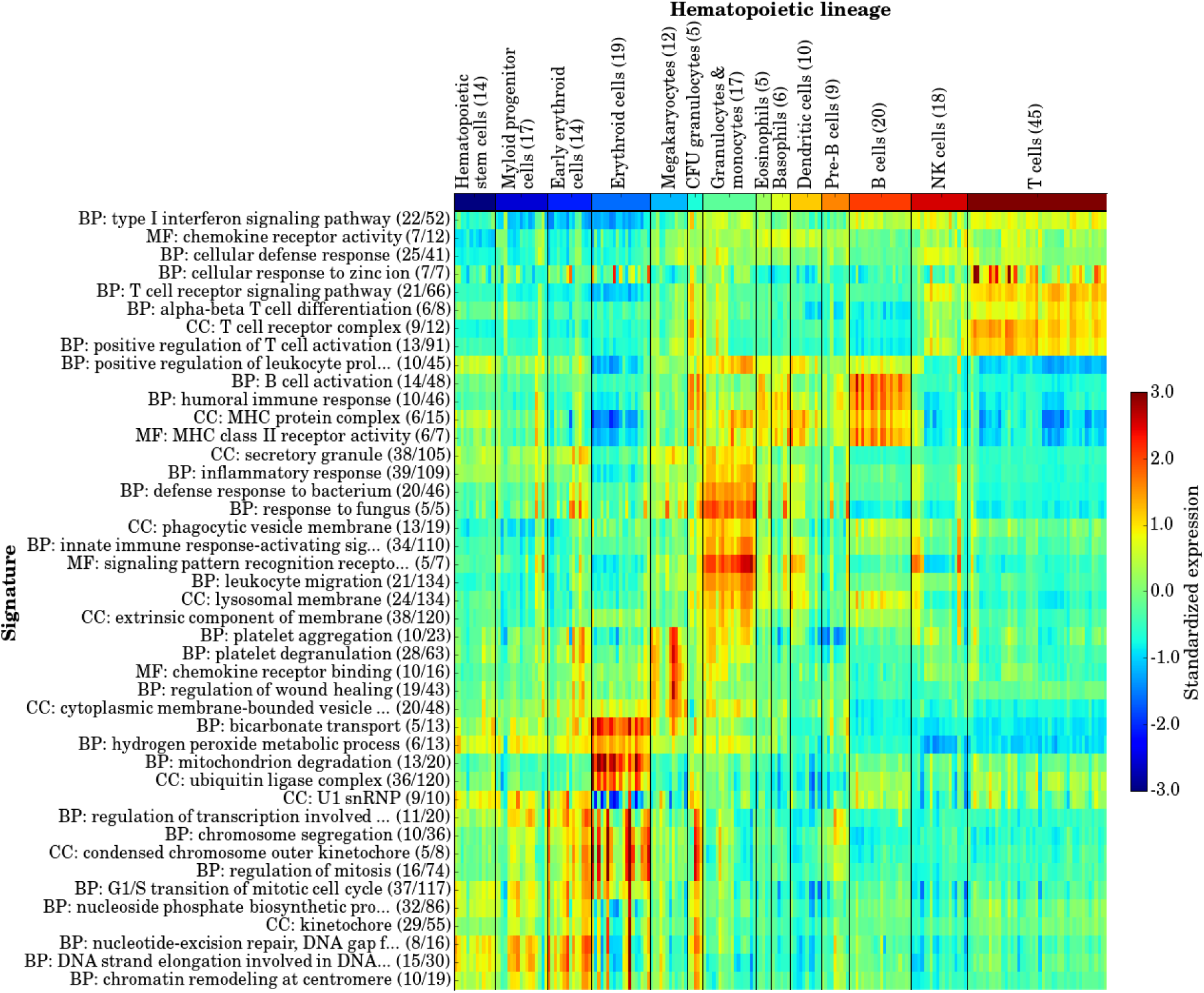
GO-PCA of 211 transcriptomes (x-axis) representing diverse hematopoietic lineages. The GO term-derived names of the 43 generated signatures are shown, with numbers in parentheses indicating the number of genes in the signature as well as the total number of genes annotated with the GO term. Lineage names are shown on top, with the number in parenthesis indicating the number of samples in each group. Lineage groupings were taken from [23].

Visual inspection of GO-PCA output immediately revealed striking associations between signatures and individual hematopoietic lineages: For example, two signatures, “bicarbonate transport” (BT) and “mitochondrion degradation” (MD) appeared strongly and exclusively associated with red blood cells. Those signatures therefore seemed to reflect known unique characteristics of these cells [24, 25], namely their ability to transport carbon dioxide and their lack of mitochondria. Further examination of these signatures revealed that the BT signature contained 5 genes (*CA1*, *CA2*, *HBA1*, *HBB*, and *RHAG*; i.e. carbonic anhydrases, hemoglobins, and a transporter) that highly contributed to PC 1, whereas the MD signature contained 13 genes that highly contributed to PC 3. The genes in these sets were associated with only 0.5% and 0.2% of the total variance in the data, respectively, highlighting the ability of GO-PCA to detect biologically relevant patterns involving only small numbers of genes against a highly heterogeneous background.

Specific associations of signatures with other cell types were also readily spotted: For example, signatures for “T cell receptor complex” (9 genes) and “positive regulation of T cell activation” (13 genes) were specifically associated with T cells. I tested whether associations between signatures and lineages were statistically significant, and found that all signatures were positively associated with at least one lineage (Figure 3), suggesting that each captured a biologically meaningful aspect of the data. Furthermore, 10 out of the 14 lineages were positively associated with at least one signature, highlighting the fact that the signatures characterized functionally diverse biological processes. The four lineages without significant associations were represented by very few samples (5-10), which likely made it more difficult for GO-PCA to identify lineage-specific patterns (they could be captured by higher principal components that were not included in the analysis), and also reduced power in detecting significant associations.

**Figure 3.**
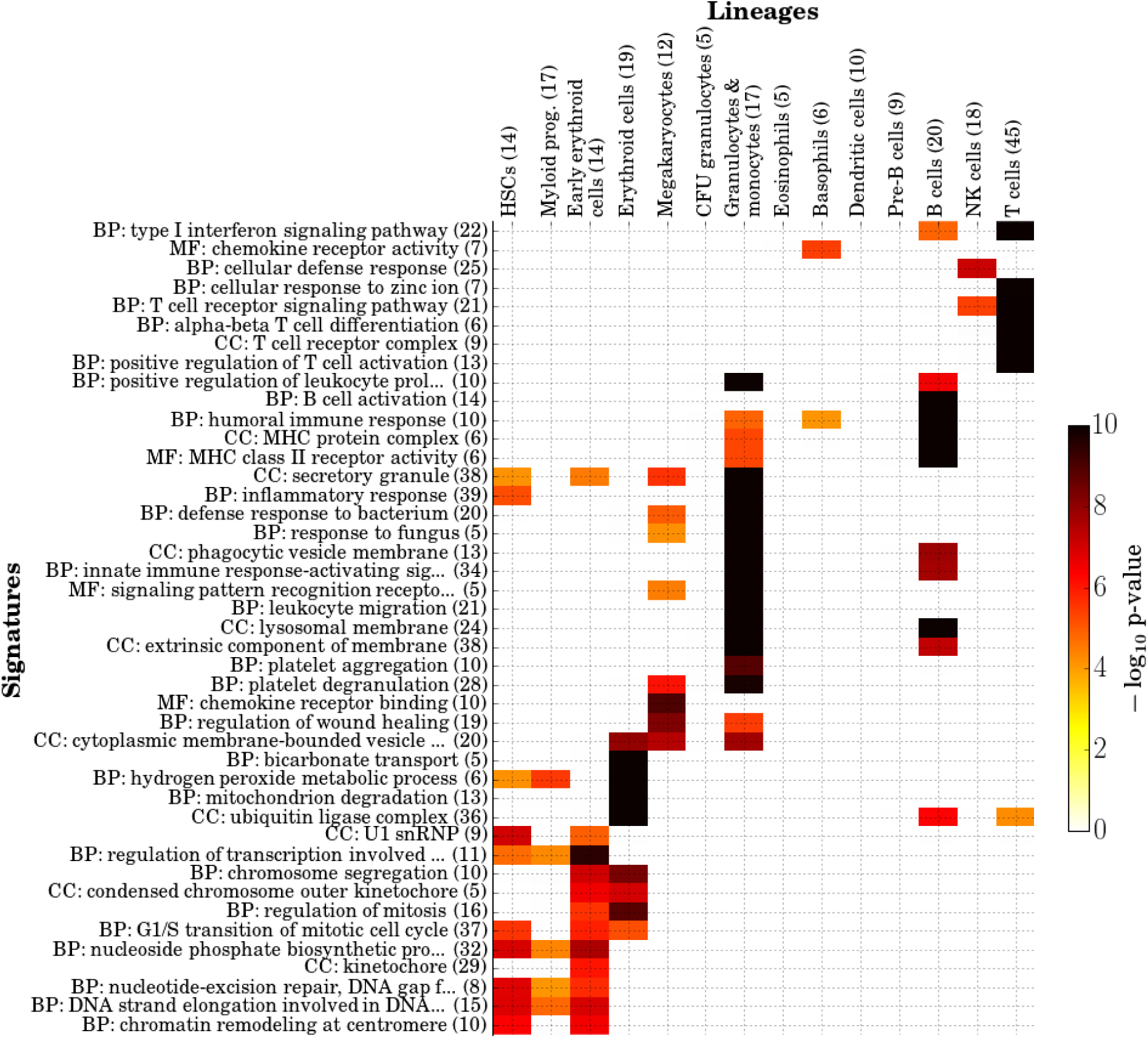
Associations between all 43 signatures discovered by GO-PCA and the 14 hematopoietic lineages represented in the data [23]. Axis labels are as in Figure 2. Shown is the significance of association, as determined by a mHG test (see Methods). Only associations with *p <* 0.05 after Bonferroni correction are shown. HSCs = hematopoietic stem cells.

On top of the many meaningful associations between signatures and lineages, some signatures also revealed significant heterogeneity among biological replicates (samples representing the same “population” within each lineage category according to the annotations provided by Novershtern et al.). This effect was observed most strongly for the cell cycle-related signatures in red blood cells, where some samples appeared much more mitotically active than others, with no obvious relationship to the three populations assayed (not shown), and depite the fact that they all shared strong expression of the BT and MD signatures. A similar effect occurred in megakaryocytes, where about half of the samples showed very strong expression of platelet-related signatures. Expression of platelet-related genes is consistent with the fact that these cells are responsible for platelet production [26], but the variation in expression had no clear relationship to the two subpopulations assayed. I suspected that these differences might be explained by batch effects, but was unable to unambiguously correlate the observed patterns with batch identifiers provided by Novershtern et al. However, the association of these patterns with specific signatures suggests that the differences represent genuine biological differences among samples, perhaps owing to dynamic nature of the differentiation processes that these cells are undergoing and the experimental protocol used to collect these cells.

### Application of GO-PCA to glioblastoma biopsies reveales subtype-specific molecular traits

After my successful application of GO-PCA to diverse cell populations from the hematopoietic system, I next asked whether GO-PCA could also facilitate exploration of heterogeneity observed in malignant disease. I therefore applied GO-PCA to 479 transcriptomes of glioblastoma biopsies from patients diagnosed with primary glioblastoma (GBM). GBMs are highly aggressive tumors, and patients have a median survival time of under 14 months [27]. Four GBM subtypes have been defined based on consensus clustering of gene expression data, and were shown to correlate with different chromosomal aberrations and mutations [28]: Classical, Mesenchymal, Proneural, and Neural. In addition, analysis of DNA methylation patterns has revealed that a subset of patients with the Proneural type exhibit a “GBM CpG island methylator phenotype” (G-CIMP), and those patients have significantly better disease outcomes [29].

GO-PCA resulted in the generation of 56 signatures, again representing divserse sources of heterogeneity (Figures S2 and S3), including what appeared to be contamination with skin cells — i.e., a small number of samples that expressed high levels of “cornified envelope” and “epidermis development” genes, and low levels of neuronal genes. Other signatures represented extracellular matrix components (e.g., collagen fibers), diverse immune system processes (e.g., a signature consisting of 20 genes annotated with “type 1 interferon signaling pathway”), as well as cell cycle-related functions and mitochondrial genes.

I systematically tested for associations between the generated signatures and the five GBM subtypes, using the classifications provided as part of the data annotations [27] (Brennan et al. treated G-CIMP as a separate subtype; see Methods). I again found strong associations between the vast majority of signatures and specific GBM subtypes (Figure 4). All five subtypes exhibited unique combinations of associated signatures, although the Classical subtpye was mainly characterized by the absence of any strong associations. The Mesenchymal subtpye showed unique associations with immune system-related processes, the Proneural subtype was uniquely characterized by high expression of cell cycle-related genes, and the Neural subtype was unique in its high expression of mitochondrial genes (suggesting elevated numbers of mitochondria in this type of tumor). However, I was most intrigued by a striking difference between the Proneural and G-CIMP subtypes, which were both grouped together in a previous expressed-based unsupervised analysis [28]. While the two subtypes shared a small number of associations (e.g., a 20-gene “somatodendritic compartment” signature), highlighting their relatedness, the G-CIMP subtype was not associated with any of the cell cycle-related signatures that were characteristic of the Proneural subtype. To verify that this was not the result of a loss of power in detecting associations for the G-CIMP subtype, which was represented by only 31 samples, I also assessed the median expression levels for each signature and subtype (Figure 4). For all cell cycle-related signatures, the G-CIMP subtype had much lower median expression than the Proneural subtype, often with levels comparable to the other subtypes. I selected the “cell cycle checkpoint” signature for further analysis, since it showed a large difference in expression and consisted of a relatively large set of genes (28/66). Its expression difference between the Proneural and G-CIMP subtypes was statistically significant (*p* = 0.008; two-sided Mann-Whitney U test), and it contained several prominent cell-cycle regulators with strong expression differences (Figure S4), including *CCNA2* (Cyclin-A2), *CCNE2* (G1/S-specific cyclin-E2) and *CDC6* (Cell division control protein 6 homolog). My results therefore suggest that lower mitotic activity is a key characteristic that distinguishes the G-CIMP subtype from its Proneural “cousin”.

**Figure 4.**
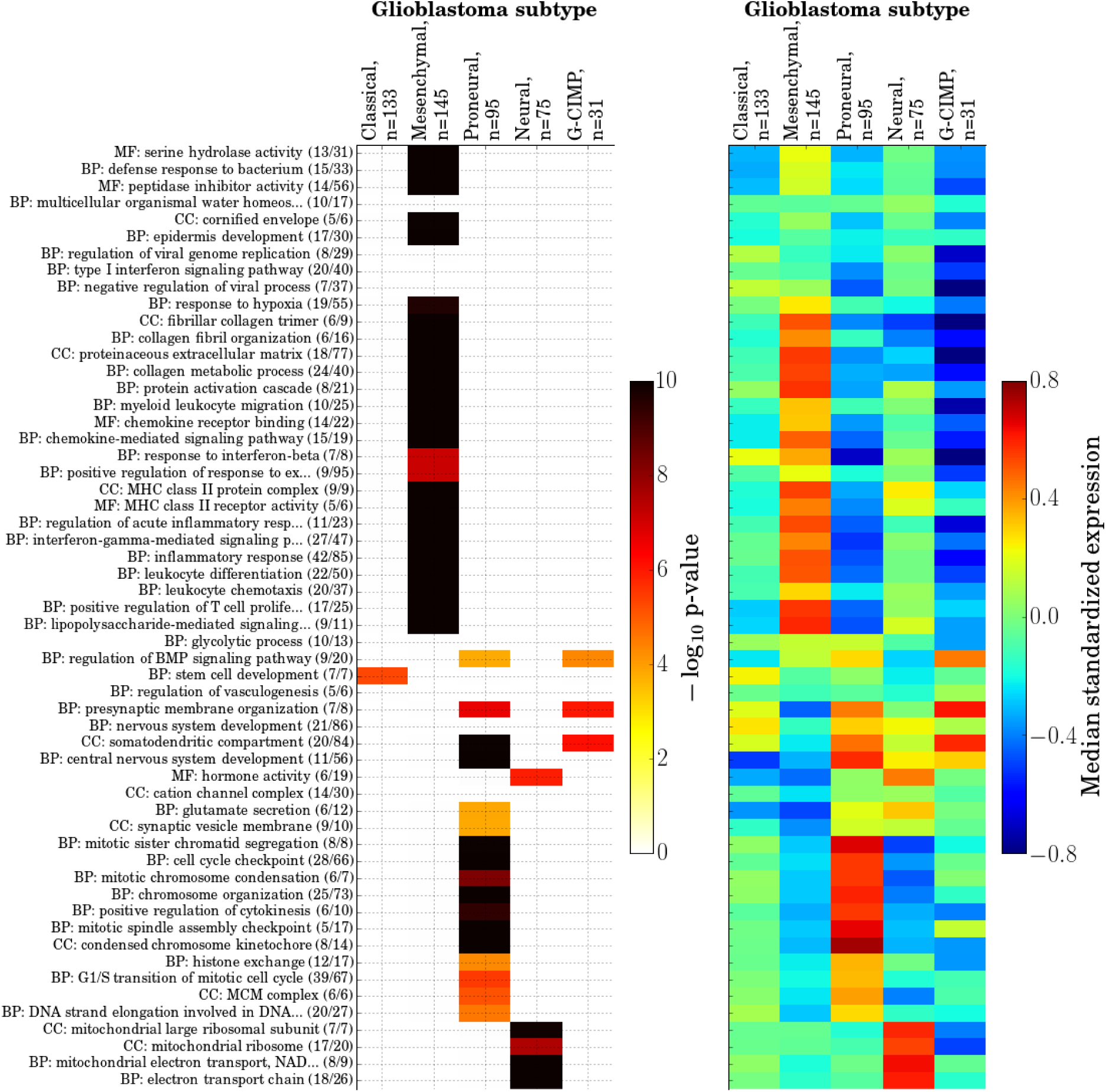
Associations between all 56 signatures discovered by GO-PCA and five previously defined subtypes of glioblastoma [27]. Left panel: significance of association, as in Figure 3. Right panel: median signature expression values for each signature and GBM subtype.

## Discussion

The high-dimensional and heterogeneous nature of transcriptomic data often makes it difficult to interpret the output of generic unsupervised algorithms, and technical biases can lead to the identification of biologically irrelevant clusters or factors [11] that further complicate the analysis. Methods incorporating prior knowledge address these challenges by biasing the analysis towards functionally coherent signals, which can improve their ability to detect relevant patterns and to provide interpretable results [12]. Here, I introduced an exploratory method that first relies on PCA to identify all major axes of variation, and then uses GO enrichment analysis as a way to focus on biologically relevant patterns. While the mHG algorithm has previously been used for GO enrichment analysis [30], GO-PCA is the first method to perform nonparametric GO enrichment analysis on genes ranked by PC loadings (to the best of my knowledge).

GO terms enriched among genes with high PC loadings provide candidate interpretations for the biological processes underlying this PC. GO-PCA takes this idea one step further, by using significant enrichments to define compact expression signatures based on the untransformed data. I believe that the signatures, as defined by GO-PCA, exhibit several properties that are desirable in an exploratory setting: Firstly, the association of each signature with a specific PC provides a way to gauge whether it is representive of a global trend in the data (associated with one of the first PCs), or a more subtle signal. Secondly, while GO-PCA’s filtering rules prioritize the terms with the strongest enrichments, they sometimes allow for the inclusion of overlapping terms (see Methods). These then offer “alternative explanations” for the biological role of the genes driving the enrichment. This limited redundancy can help reduce the likelihood of drawing wrong conclusions based on the enrichment of a single term that might not accurately reflect the underlying biological process. Thirdly, the number of genes that contribute to a given signature is relatively small (typically 10-20, and not more than 50). In combination with the fact that the signature value represents the unweighted average expression of its genes (after standardization), this provides an immediate ability to understand how signature values are related to the underlying raw data, which would not be the case for signatures that are defined using arbitrary linear combinations or more complex functions. The averaging also imparts the signature with a certain level of robustness against measurement errors. Finally, the signatures offer a powerful opportunity to assess the relationship between different biological processes (demonstrated here in Figures S1 and S3).

Prior knowledge can be seen as a bias that is introduced to the unsupervised analysis. Some methods that incorporate prior knowledge enable control over the strength of this bias using an explicit tradeoff parameter [12]. GO-PCA is not designed as a flexible-tradeoff method. While its first step, PCA, does not incorporate any prior knowledge, the GO enrichment testing and signature generation steps use prior knowledge to identify compact sets of genes that are both correlated and functionally related. As a result, GO-PCA excludes the measurements of the majority of genes from the final output. In my view, the introduction of strong biases are easily justified by the large gains in expressiveness and interpretability associated with the signatures generated. While some axes of variation might be missed due to incomplete gene annotations, I feel that the diversity of the processes identified in my analyses is a testament to the comprehensive nature of the human UniProt-GOA annotations [21].

I did not explore applications to datasets from other organisms, and in both use cases demonstrated here, the transcriptomic data were generated using microarrays. However, I have good reason to believe that in principle, GO-PCA will also perform well for data from other organisms and generated using different platforms (e.g., RNA-Seq). GO-PCA relies on two nonparametric methodologies (PCA and mHG) and therefore avoids strong assumptions about distributional properties of the data. Thus, generally speaking, GO-PCA will be applicable whenever PCA can be expected to recover meaningful axes of variation. Its performance obviously depends on the quality of the gene annotations available, but this is true of any annotation-driven approach. The only parameter that might require careful consideration on a case-by-case basis is *D*, the number of principal components tested. The more components tested, the more sensitive the methods becomes with respect to detecting weak patterns, potentially associated with only a small number of genes and/or samples. However, this comes with the tradeoff of more false positive enrichments due to multiple testing (if the p-value threshold used in the enrichment analysis is not adjusted by the number of PCs tested), or reduced power in the enrichment analysis (if it is). Rather than testing a larger number of principal components, I would advocate careful study of the main sources of heterogeneity in the data, estimating and possibly removing their contributions using appropriate methods, and then re-running GO-PCA.

In future work, I will consider statistical approaches to selecting the number of PCs to test, rather than using an arbitrary variance threshold. Specifically, it has recently been suggested to assess the association between individual variables (genes) and principal components using a “jackstraw” approach [31]. Using this methodology, GO-PCA could decide to stop testing the next PC when there are no (or not enough) genes significantly associated with it. Moreover, it could use this information to rank genes based on significance of association (instead of using the loadings directly), and flexibly adjust *L* for each PC in the mHG algorithm, e.g. based on an FDR criterion [32]. This could further improve the power of detecting enriched terms. On an Intel(R) Xeon(R) E5620 CPU, it took GO-PCA 452s to test GO enrichment for 12 PCs of the hematopoietic dataset (38s/PC), and 436s to test GO enrichment for 15 PCs of the glioblastoma dataset (29s/PC). The algorithm required roughly 1 GB of memory. In order to further improve runtime, the algorithm could easily be parallelized, since GO enrichment can be tested for each PC independently. In conclusion, the methodology described here provides a powerful and versatile framework for the exploration of gene expression data, and demonstrates the potential of unsupervised algorithms that directly incorporate prior knowledge.

## Methods

### Obtaining a list of all protein-coding genes

GENCODE Version 19 gene annotations were downloaded from http://www.gencodegenes.org and filtered for transcripts that had a “transcript_type” attribute set to “protein coding”. The set of all associated “gene name” attribute values was taken as the list of all protein-coding genes, yielding 20,114 genes.

### Obtaining the GO ontology and GO gene annotations

The Gene Ontology structure (`go-basic.obo`) and UniProt Gene Ontology Annotations (`gene_association.goa_human.gz`) were downloaded from http://geneontology.org on 01/18/2015. All annotations were propagated up the GO graph based on the “is_a” relationships (i.e., a gene that is annotated with a particular term was also annotated with all parent terms, since those represent more general categories). For GO terms in the “cellular component” domain, “part_of” relationships were treated the same as “is_a” relationships. All annotations with evidence code “IEA” (inferred from electronic annotation) were ignored, in order to restrict annotations to manually curated terms. 43% of all UniProt-GO annotations fell into this category. I further removed GO terms that were either too broad (defined as having more than 200 genes annotated with them), or too specific (defined as having less than 10 genes annotated with them). Finally, I identified all instances where two or more GO terms had identical sets of genes annotated with them. For example, the term “organellar large ribosomal subunit” (GO:0000315) had the same 21 genes annotated with it as its child term, “mitochondrial large ribosomal subunit” (GO:0005762). In cases like this, where a parent term had identical sets of annotated genes as the child term, I removed the parent term, resulting in the exclusion of 280 terms. This resulted in a final set of 4,418 terms that were neither too broad nor too specific, and not redundant with any related term.

### Principal component analysis (PCA)

PCA was performed using the `decomposition.PCA` module from the scikit-learn Python package [33] (http://scikit-learn.org).

### Nonparametric GO enrichment analysis using the minimum hypergeometric (mHG) test

Given a ranked list of protein-coding genes (where the ranking was defined by the loadings associated with a particular principal component), I tested for GO enrichment using a Cython [34] (http://cython.org) implementation of the minimum hypergeometric (mHG) statistic [22]. The mHG statistic calculates a hypergeometric enrichment p-value (equivalent to Fisher’s exact test) for all *N* possible thresholds in a ranked list of *N* binary variables, and then selects the threshold associated with the best (lowest) p-value. Due to the many tests performed, this p-value cannot be taken at face value and should be thought of as an enrichment statistic. I refer to this value as *s*^mHG^. The mHG test then employs a dynamic programming algorithm [22] to calculate the expected probability *p*^mHG^ for obtaining a statistic as small as or smaller than *s*^mHG^, when given a random permutation of the ranked list. This algorithm has a time complexity of 𝒪(*N*^2^), as opposed to the computationally infeasible 𝒪(*N*!) time complexity that would be required for explicitly enumerating all possible permutations. By definition, *p*^mHG^ is the exact p-value associated with *s*^mHG^.

I furthermore extended the mHG test to allow the user to limit the thresholds tested based on two criteria, associated with two “hyperparameters” that I refer to as *X* and *L*. I therefore refer to this extended version of the mHG as XL-mHG. The first criterion ignores all thresholds at which less than *X* positive (i.e., 1-valued) variables have been encountered. This criterion is designed to improve the test’s robustness: For example, if there are highly correlated variables, a small number of positive examples at the top of the list might result in spurious enrichment. This can be avoided by setting *X* to a value deemed large enough to guard against such cases. In GO-PCA, I set *X* = 5 (note that setting *X* to a relatively large value like 20 would result in a very robust test, but it would also make it impossible to detect enrichment for a significant number of GO terms which have fewer than 20 genes annotated with them). The second criterion limits the thresholds tested to the first *L* ranks (*L < N*). This criterion is designed to avoid cases where a very minor enrichment (e.g. 1.2-fold) is reported as highly significant, solely because it is observed at a very low threshold. For example, in a list of 10,000 genes, testing thresholds of 5,000 and lower often makes no sense, as any biologically meaningful enrichment is expected to result from gene ranked much higher in the list. In my experience, any enrichment that can only be detected at such a low threshold is likely the result of technical biases and does not constitute a biologically meaningful enrichment. (This effect was also recognized as an important problem in the development of GSEA [13]). In GO-PCA, I use *L ≈ N/*8, due to the “two-sided” nature of the test (for a “one-sided” test, I would use *L ≈ N/*4).

## GO-PCA

The “backbone” of GO-PCA consists of a simple algorithm: 1) Perform PCA on the gene expression matrix (treating the genes as variables and the samples as observations), and extract the gene loadings associated with the first *D* principal components. I determine *D* by requiring that any PC analyzed should capture at least 1% of the total variance. Since PCs are sorted based on their % variance explained in decreasing order, this determines a unique *D*. 2) For each PC, rank genes by their loadings, in both ascending and descending order (this produces two gene rankings). 3) For each ranking, determine enriched GO terms using the XL-mHG test and a bonferroni-corrected p-value threshold. I use a threshold of *α* = 0.01 and a bonferroni correction for the the number of tests performed for each PC. I test *m* = 4418 GO terms for both rankings, so the correction factor is 2\**m ≈* 10, 000, and the corrected p-value threshold is *≈* 1 * 10^−6^. 4) For each enriched GO term, use the minimum hypergeometric threshold determined by the XL-mHG test, and find all genes annotated with the GO term that are above the threshold. For each of those genes, standardize its expression (substract the sample mean and divide by the sample standard deviation), and then average the standardized expression values across all those genes. This average is the signature expression value. Label the signature with the name of the GO term.

Along with this “backbone”, GO-PCA also filters the GO terms found to be enriched at each step to avoid redundancies that result from the nested structure of GO terms. This “local” filter is applied independently for each PC and each gene ranking (i.e., it is applied twice per PC, once for each ranking). The key intuition that I used to devise these filters is that among the significantly enriched GO terms, some GO terms are more enriched than others, and that this redundancy can be tested by asking whether the enrichment of a specific GO term is still significant after removing genes that were associated with more enriched terms. To obtain an estimate of the enrichment effect size, I again used the threshold found by the XL-mHG algorithm to calculate a fold enrichment value for each enriched GO term (defined as the observed number of genes annotated with the term that fall above the threshold, divided by the expected number). I then ranked enriched GO terms by their fold enrichment (in descending order), and applied the following filtering procedure: 1) Initialize a set of “seen” genes with the genes in the signature derived from the first GO term. 2) Remove all “seen” genes from the data and re-test the enrichment of the second term using the XL-mHG algorithm. If the test is still significant (using the same p-value threshold as used before), keep the signature derived from the second term and add its genes to the set of “seen” genes. If the test is not significant anymore, discard the signature associated with the second term. 3) Repeat step 2 for the third term, then the fourth, and so on, until all enriched GO terms have been tested for redundancy.

While the previously described filter helps avoid redundancies within each principal component, I also found that strong biological effects (e.g., differences in the expression of cell cycle genes) are sometimes associated with multiple principal components. To mitigate these cross-PC redundancies, I also applied a second “global” filter that removes a signature generated by a PC if its associated GO term (or one of its parents) was previously used to generate a signature for another PC. (Note that this implies that enrichments associated with earlier PC are prioritized over enrichments associated with later PCs, motivated by the fact that the earlier PC captures a larger fraction of the total variance). In summary, the “local” and “global” filters signifincantly limit the redundancy of signatures. However, when presented with “sufficient evidence”, they allow signatures from the same PC to exhibit significant overlap.

### Analysis of hematapoeitic expression data

I downloaded the hematopoietic dataset generated by Novershtern et al. [23], including sample annotations, from http://www.broadinstitute.org/dmap/home. This data is fully processed, and contains only genes that were expressed in the majority of samples from one cell type (see [23] for details). Out of the 8,968 genes in the data, I excluded 473 (5.2%) that were not in my list of protein-coding genes (see above). For visualization and analysis, I used the annotations provided in the file `DMap_sample_info.022011.xls` to sort sorted samples based first on lineage (column 6), then on population (column 5), and then on batch (column 4). For the XL-mHG algorithm, I set X=5 and L=1,000. Signatures were sorted using the leaf ordering of a dendrogram generated by hierarchical clustering with pearson correlation as the distance metric and using average linkage, as implemented in scipy’s `clustering.hierarchy.linkage` function.

### Analysis of glioblastoma expression data

I curated a set of 479 glioblastoma transcriptomes from TCGA based on clinical annotation data from Supplementary Table 7 (“Clinical and Molecular Subclass Data Table”) in [27]. I first excluded patients that did not present with primary GBM (i.e., I excluded patients if they did not have “NO” in Column 2, “Secondary or Recurrant”), retaining 516 patients. I then downloaded annotation data for all GBM datasets in the TCGA data freeze from Oct 10, 2012 (`data.freeze.txt` from https://tcga-data.nci.nih.gov/docs/publications/gbm_2013/), and filtered for rows with column 5/“DATATYPE” equal to “Expression-Gene” and column 8/“DATA LEVEL” equal to “3”). I then used these additional annotations to exclude samples that were annotated with any of the following terms (column 13, “ANNOTATION CATEGORIES”): “item in special subset”, “normal class but appears diseased”, “qualified in error”. This resulted in the exclusion of 37 samples, and a final dataset of 479 samples. The expression data for these patients was exctracted from the “Level 3” expression data (`GBM.Gene_Expression.Level_3.tar` from https://tcga-data.nci.nih.gov/docs/publications/gbm_2013/). Of the 12,042 genes contained in those data, 1,220 (10.1%) were not recognized as canonical protein-coding gene names (see above) and thus excluded from the analyses.

Instead of treating the remaining 10,822 protein-coding genes as the “universe” of genes in the GO enrichment analysis performed as part of GO-PCA, I performed variance filtering [35] and retained only the top 5,500 most variable genes (removing *≈* 15% of the total variance in the data). The main motivation for this is to guard against artifacts in the GO enrichment analysis, which could result as a consequence of the fact that glial cells, and human cells generally, only express a subset of all genes in the human genome. The most variable genes along any given PC therefore tend to be “enriched” for glial-specific genes whenever all human genes (or an unbiased subset — i.e., the genes represented on the array) are treated as the universe. By only retaining the *≈* 50% most variable genes, the universe is more similar to the set of genes with expression in glial cells. For the XL-mHG algorithm, I set X=5 and L=750. Signatures were sorted using hirarchical clustering, as in the anlaysis of the hematopoietic dataset.

### Testing for association between signatures and subsets of samples

I used the XL-mHG test in order to test for association between individual signatures and subgroups of samples in the following way: For each subgroup, all samples in the subgroup were labeled “1”, all others “0”. Then, for each signature, the entire set of samples was ranked in descending order based on their signature values, and mHG was used to test for enrichment of the subgroup samples at the top of the list. For an analysis of *d* signatures and *m* subgroups, *d* * *m* tests were performed, and the bonferroni-corrected p-value threshold used to call significant associations was therefore calculated as 0.05*/*(*d* * *m*). The parameters used for mHG-XL were X=3 / L=100 for the hematopoietic data, and X=10 / L=200 for the glioblastoma data.

### Software

GO-PCA is free software and can be found at https://github.com/flo-compbio/gopca.

## Ackowledgements

I would like to thank Dr. Sandeep Dave for his support, Dr. Anupama Reddy and Dr. Jyotishka Datta for helpful discussions, as well as Dr. Anupama Reddy and Dr. Alexander Hartemink for their feedback on the manuscript. Furthermore, I would like to thank Dr. Zohar Yakhini for introducing me to the mHG algorithm.

## Supplemental Figures

**Figure S1.**
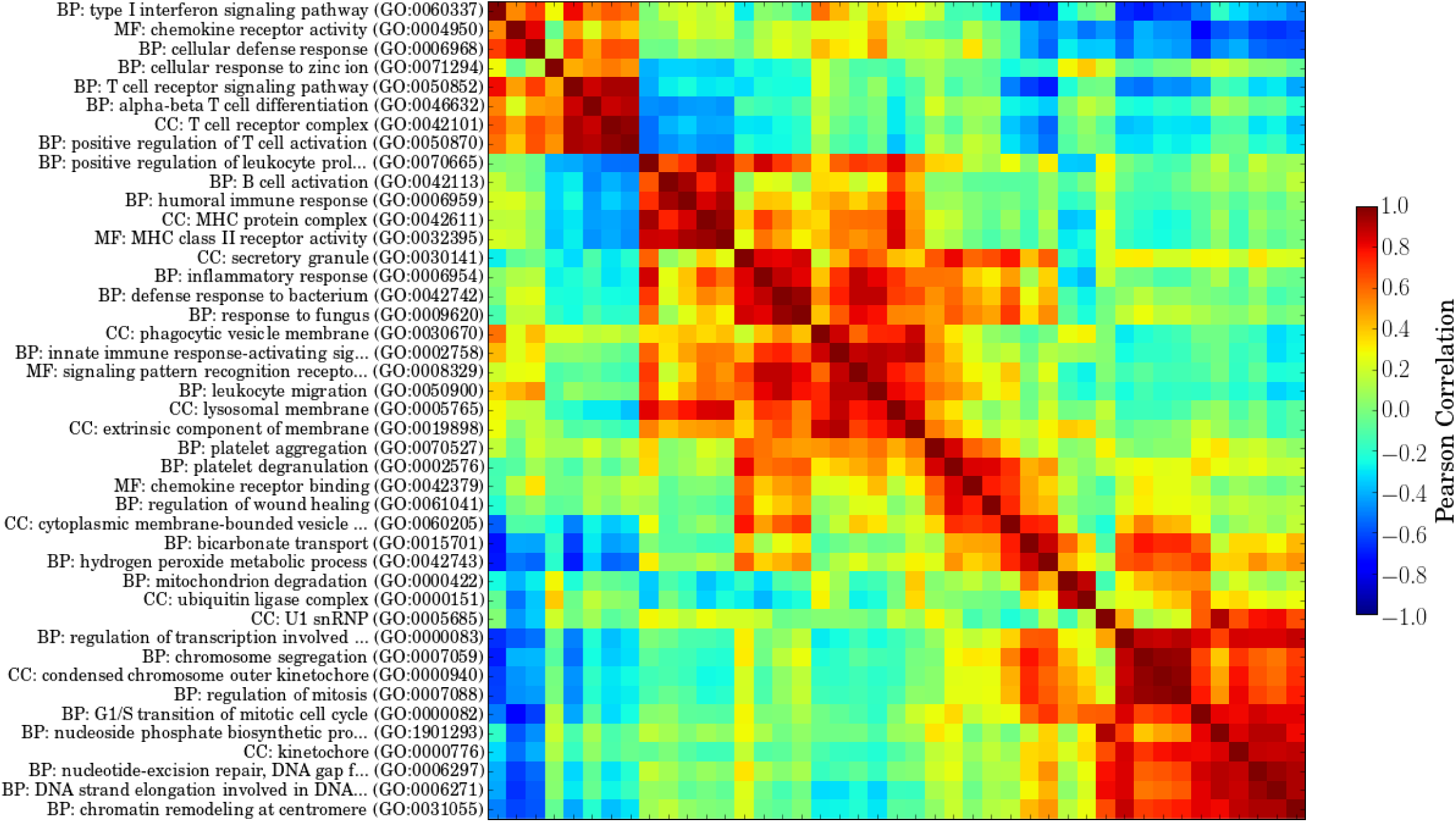
Heatmap showing pair-wise correlations among all 43 signatures generated by GO-PCA analysis of 211 transcriptomes representing cell populations from diverse hematopoietic lineages [23].

**Figure S2.**
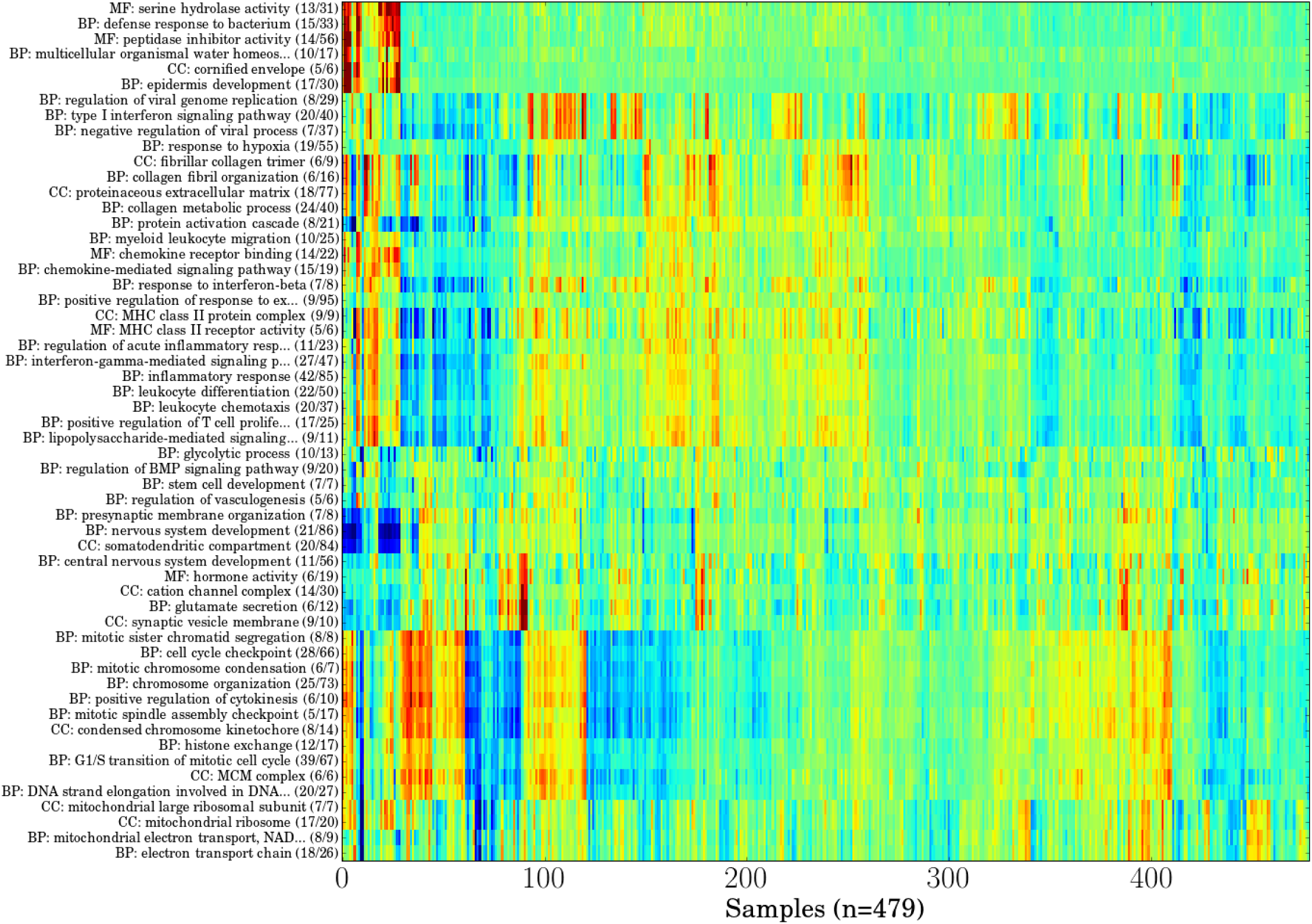
Heatmap showing all 56 signatures generated by GO-PCA analysis of biopsies from 479 patients newly diagnosed with glioblastoma [27]. Both samples (x-axis) and signatures (y-axis) are ordered using hierarchical clustering. For explanation of axis labels and color scheme, see Figure 2.

**Figure S3.**
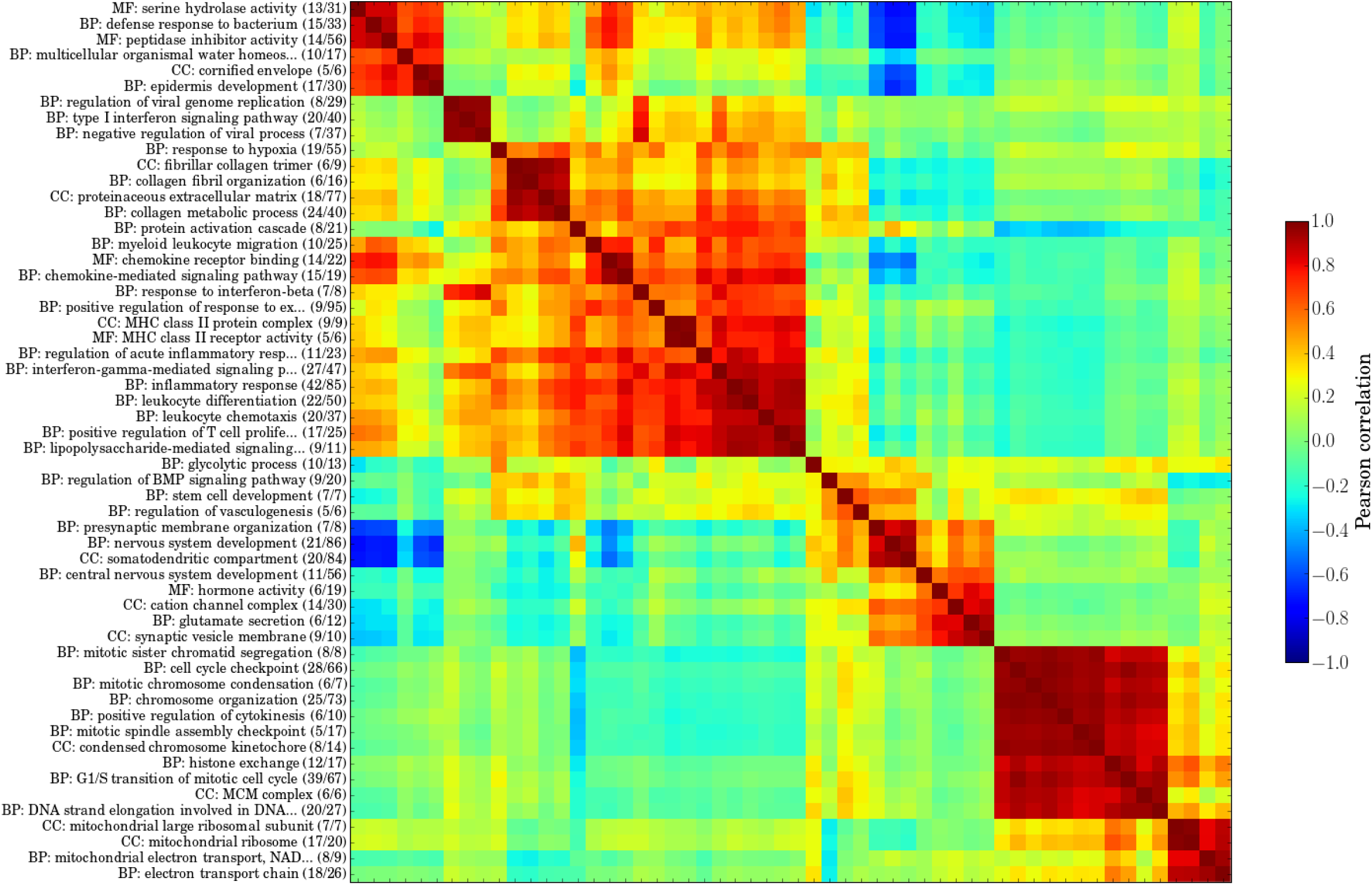
Heatmap showing pair-wise correlations among all 56 signatures generated by GO-PCA analysis of biopsies from 479 patients with primary glioblastoma [27] (cf. Figure S2).

**Figure S4.**
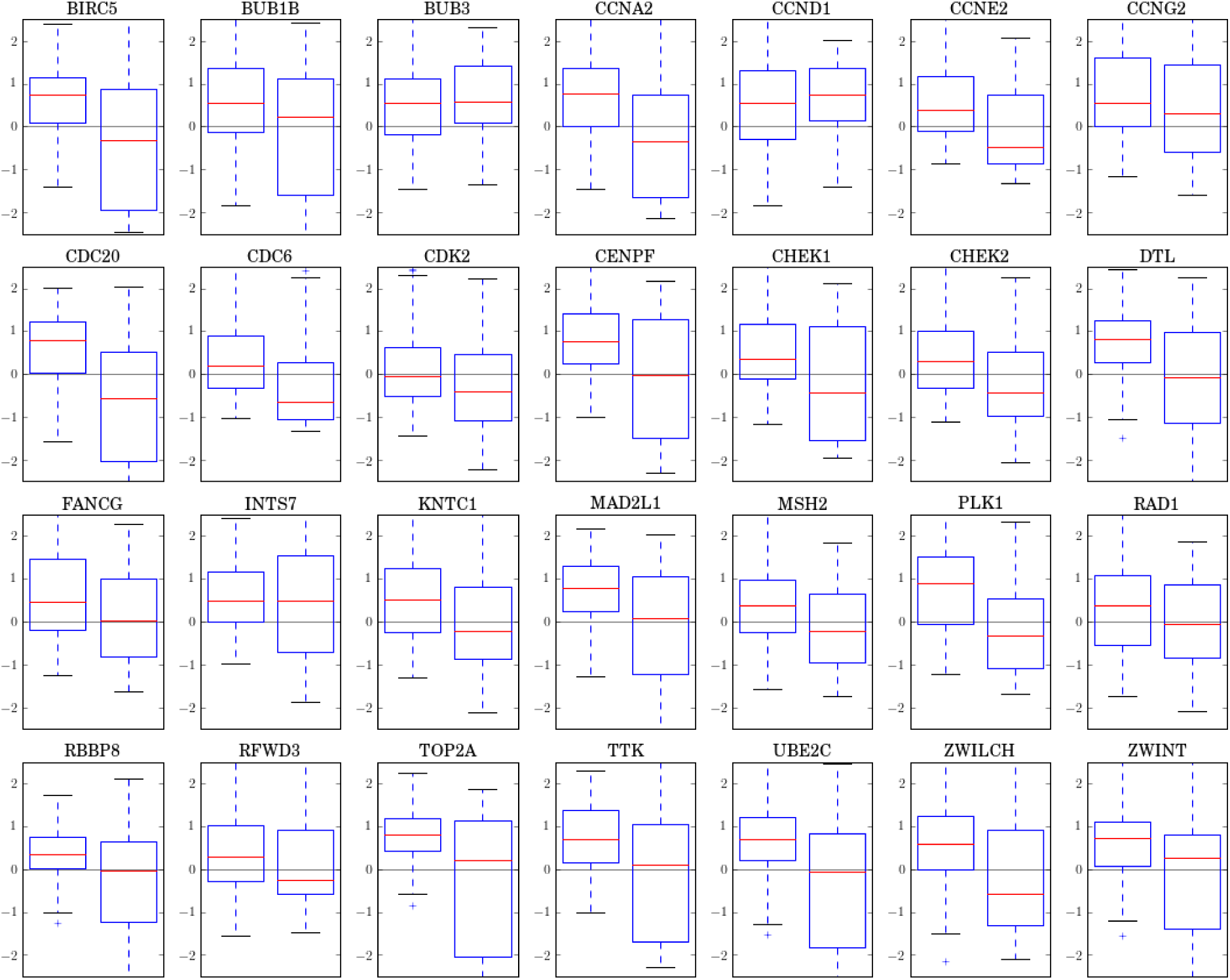
Boxplots showing expresison differences between the Preneural (left) and G-CIMP (right) subtypes for all 28 genes in the “cell cycle checkpoint” signature. The expression levels for each gene are standardized (i.e., mean = 0, and standard deviation = 1), based on data from the entire GBM dataset.

